# Exogenously driven neural reactivation of spatially matching visual working-memory contents

**DOI:** 10.64898/2026.01.22.701110

**Authors:** Águeda Fuentes-Guerra, Elisa Martín-Arévalo, Freek van Ede, Carlos González-García

## Abstract

Selective attention is often divided into voluntary (goal-directed) and involuntary (stimulus-driven) forms, a distinction extensively studied for attention to external sensory input. In contrast, internal selective attention—directed toward representations held in working memory (WM)—has been considered primarily for voluntary influences. Recent behavioral evidence suggests that task-irrelevant external stimuli can also influence internal selection of feature-matching WM representations involuntarily, yet the neural mechanisms underlying these effects remain unclear. Here, we tested whether an uninformative exogenous spatial retro-cue presented during a WM delay can act as a selective “ping” and reactivate spatially matching WM content at the level of its representational category. Male and female human participants memorized complex contents presented at distinct locations, followed by unpredictive and task-irrelevant spatial retro-cues that conveyed no category information. Using temporally resolved multivariate EEG decoding, we observed category-specific reactivation of spatially matching WM representations following these cues, providing direct neural evidence for stimulus-driven, involuntary attentional selection within WM, ahead of the memory test. Moreover, neural responses to the subsequent memory-probe were also modulated by cue congruency, consistent with the notion that exogenous influences begin early during sensory processing while also shaping later decision-related processes. Finally, drift-diffusion model analyses revealed that this involuntary cueing effect was primarily driven by increased evidence accumulation. Together, these findings illuminate the mechanisms by which external events can automatically and involuntarily penetrate the internal cognitive workspace.

**Significance Statement:** The processes underlying involuntary attentional selection of internal information held in our working-memory system have been largely overlooked. Our study shows that external stimuli can automatically select information that is temporarily held in our mind. We found that uninformative, task-irrelevant visual cues can selectively “ping” and reactivate specific working-memory contents, despite carrying only spatial information. We demonstrate that these cues trigger category-specific neural reactivation and shape later decision-making by accelerating evidence accumulation. This extends current knowledge on how the outside world can transiently interact with our internal cognitive workspace.

## Introduction

Selective attention is commonly categorized into voluntary (goal-directed) and involuntary (stimulus-driven) (Jonides, 1981; Corbetta et al., 2008). While this distinction has been extensively studied in the context of external selective attention to sensory input (Carrasco, 2011; Chica et al., 2013; Chica et al., 2014), internal selective attention—attention directed toward specific sensory representations held in working-memory (WM) (Chun et al., 2011; van Ede & Nobre, 2023), has been predominantly examined as a voluntary process (Gazzaley & Nobre, 2012; Gunseli et al., 2015; Gunseli et al., 2019; Griffin & Nobre, 2003; Landman et al., 2003; Rerko et al., 2014; Souza et al., 2014, 2015).

Recent evidence suggests that task-irrelevant external stimuli can also involuntarily influence internal selection by facilitating feature-matching WM representations (Cipriani et al., 2024; Ding et al., 2024; Fuentes-Guerra et al., 2025a, Fuentes-Guerra et al., 2025b; Han et al., 2023; Joe & Kim, 2023; Liao et al., 2025; van Ede et al., 2020; for similar findings in iconic memory, see also Sergeant et al., 2013). However, the neural mechanisms underlying these exogenous influences remain poorly understood. Behavioral studies have shown that non-predictive retro-cues (i.e., cues presented between the offset of a WM array and the onset of a WM probe) can enhance performance for spatially matching representations (Joe & Kim, 2023; Cipriani et al., 2024; Fuentes-Guerra et al., 2025a, Fuentes-Guerra et al., 2025b). These effects appear to arise from both enhanced efficiency in extracting relevant information for subsequent decisions and reduced time to access relevant memory representations upon probe presentation (Cipriani et al., 2024; Fuentes-Guerra et al., 2025a, Fuentes-Guerra et al., 2025c). Beyond behavioral evidence, other studies have examined indirect spatial markers of internal attention (Ding et al., 2024; van Ede et al., 2020) through biases in fixational gaze behavior, providing more direct support for an involuntary “retro-capture” effect whereby external stimuli trigger the selection of feature-matching internal representations.

While there is behavioral and gaze evidence that exogenous non-predictive retro-cues can select and prioritize WM contents, a critical and open question remains: whether and how unpredictive and task-irrelevant sensory inputs can reactivate matching memories at the level of their representational content. Concurrently, studies using sensory “pinging” have shown that irrelevant inputs can revive latent WM traces (Barbosa et al., 2021; Karabay et al., 2025; Wolff et al., 2015, 2017; Yang et al., 2025; Zhang & Luo, 2023). However, to date, such effects have been demonstrated as global reactivations, without evidence of content-specificity. Here, we hypothesized that a non-predictive exogenous spatial retro-cue could act as a selective “ping” and automatically reactivate the spatially matching selected content. Note how such a mechanism would resemble the phenomenon of targeted memory reactivation, an established framework in long-term memory and sleep research (Rasch & Born, 2007; Lewis & Bendor, 2019; Herszage & Censor, 2017), where cues associated with prior learning during sleep or rest can selectively strengthen corresponding memory traces. We thus hypothesize that similar principles may operate during maintenance in WM, which remains underexplored.

The current study bridges these previously independent lines of research by testing whether an uninformative exogenous retro-cue presented during a WM delay can act as a selective “ping” of spatially matching WM content by reactivating its specific representation. Using temporally resolved multivariate decoding of electroencephalography (EEG) data, we examined whether such task-irrelevant and uninformative spatial retro-cues elicit category-specific reactivation of spatially matching WM representations. Specifically, we tested whether the animacy category of previously memorized items could be decoded from the EEG signal following spatial retro-cues that themselves conveyed no category-level information but merely matched the location of previously encoded animate or inanimate memory content.

To anticipate our results, we provide direct neural evidence for stimulus-driven, involuntary attentional selection and associated content re-activation in WM. In addition, we show how the initial and later processing of subsequent probes is contingent on the congruency of preceding exogenous retro-cues, corroborating that exogenous cueing effects during WM start ahead of the probe and do not exclusively affect later processes associated with decision-making and response execution.

## Materials and Methods

### Data and Code Availability

Raw data, task design scripts and analysis’ scripts for this experiment can be found at (https://openneuro.org/datasets/ds007180) and (https://osf.io/e3gxj/overview?view_only=ad6c91039e5f40bc9bebd8d998a27812), respectively.

### Methods

#### Participants

Thirty-five naïve volunteers participated in the experiment. Ten participants were excluded from the final sample due to technical issues with the triggers (n=1), excessive noise in the EEG signal (n=1), or behavioral performance below 60% accuracy (n=8). The final sample consisted of 25 participants (7 cis-males, 18 cis-females, mean age of 23 years, SD=3.15), similar to previous studies using related paradigms and outcome measures (van Ede et al., 2017; Liu et al., 2022). Participants were recruited through the Brain, Mind, and Behavior Research Center (CIMCYC) experimental participation website. Eligibility criteria included being between 18 and 35 years old, having normal or corrected-to-normal vision, and providing written informed consent. Participants received monetary compensation (€5 per 30 minutes) upon completion of the experiment.

The study was conducted in accordance with the ethical guidelines of the University of Granada and adhered to the principles of the Declaration of Helsinki (2013 revision, Brazil). The protocol was approved by the University of Granada Ethics Committee (Ref: 1816/CEIH/2020).

#### Apparatus, stimuli and procedure

The experiment was run on a computer equipped with an Intel Core i7-3770 CPU @ 3.40 GHz x8, connected to a 24-inch Benq XL2411T monitor (1920 × 1080 resolution, 350 cd/m^2^ brightness). Participants were seated approximately 65 cm from the screen in a dimly lit room. Stimulus presentation and data collection were controlled using PsychoPy v2021.2.3.

Each trial began with an encoding display consisting of a central fixation point and two placeholder boxes (200 × 200 pixels, with a 10-pixel border), positioned to the left (x = –250, y = 75) and right (x = 250, y = 75) of fixation. Each placeholder contained a grayscale image (200 × 200 pixels) of either an animate (non-human animals) or inanimate object (vehicles, tools, house appliances, etc.). Images were sourced from publicly available databases (Brady et al., 2008, Brady et al., 2013; Brodeur et al., 2014; Griffin et al., 2022; Konkle et al., 2010; Hebart et al., 2019), resulting in a pool of 2580 unique images (1290 animate, 1290 inanimate). All images were preprocessed to remove backgrounds, center the object, and convert to grayscale to enhance perceptual distinctiveness.

Peripheral retro-cues were implemented by increasing the border thickness of the placeholder from 10 to 30 pixels. The task was a choice-reaction time task embedded in an exogenous retro-cueing paradigm with a cue-target onset asynchrony (CTOA) of 250–350 ms, previously shown to elicit reliable cueing effects (Martín-Arévalo et al., 2013, 2016, Martín-Arévalo et al., 2021; Fuentes-Guerra et al., 2025a).

Each trial proceeded as follows (see Fig. 1): the encoding display (fixation, placeholders, and two novel images) was presented for 1,000 ms. One image appeared on an orange background and the other on a green background. Participants were instructed to form stimulus–response (S–R) associations based on background color. For example, orange stimuli were to be associated with bimanual index finger responses, and green stimuli with bimanual middle finger responses. The color–response mapping was counterbalanced across participants and remained constant throughout the experiment. S-R mappings were orthogonal to stimulus location. Importantly, each stimulus pair was trial-unique and never repeated. Following the encoding display, a 500 ms delay screen (placeholders and fixation only) was shown (see Souza & Oberauer, 2016). A non-predictive peripheral retro-cue (validity of 50%) then appeared for 50 ms on one of the two placeholders. Participants were instructed to neglect the task-irrelevant spatial retro-cue and maintain central fixation. After a jittered fixation interval (200–300 ms), a centrally presented black and white target image (x = 0, y = 75) appeared for 1,200 ms. Participants responded using both hands: pressing “S” and “L” simultaneously for middle finger responses, or “D” and “K” for index finger responses. Both keys had to be pressed simultaneously for the response to be considered correct; partial or mixed responses were marked as incorrect. Reaction times (RTs) were recorded for both fingers, but only the fastest RTs were used for analysis. In a given trial, one of the encoding images was always ‘animate’ while the other was ‘inanimate’. Stimulus animacy was never task-relevant. The retro-cue equally often matched the location of the previous animate or the inanimate image. In 3% of trials, a novel image not shown during encoding was presented as the target (“catch trials”). Participants were instructed to press the spacebar with their thumbs in these cases. Catch trials were included to discourage strategies that reduced WM load (e.g., encoding only one item). The inter-trial interval, in which the screen remained empty, had a jittered duration between 1,000–1,500 ms.

**Fig. 1.**
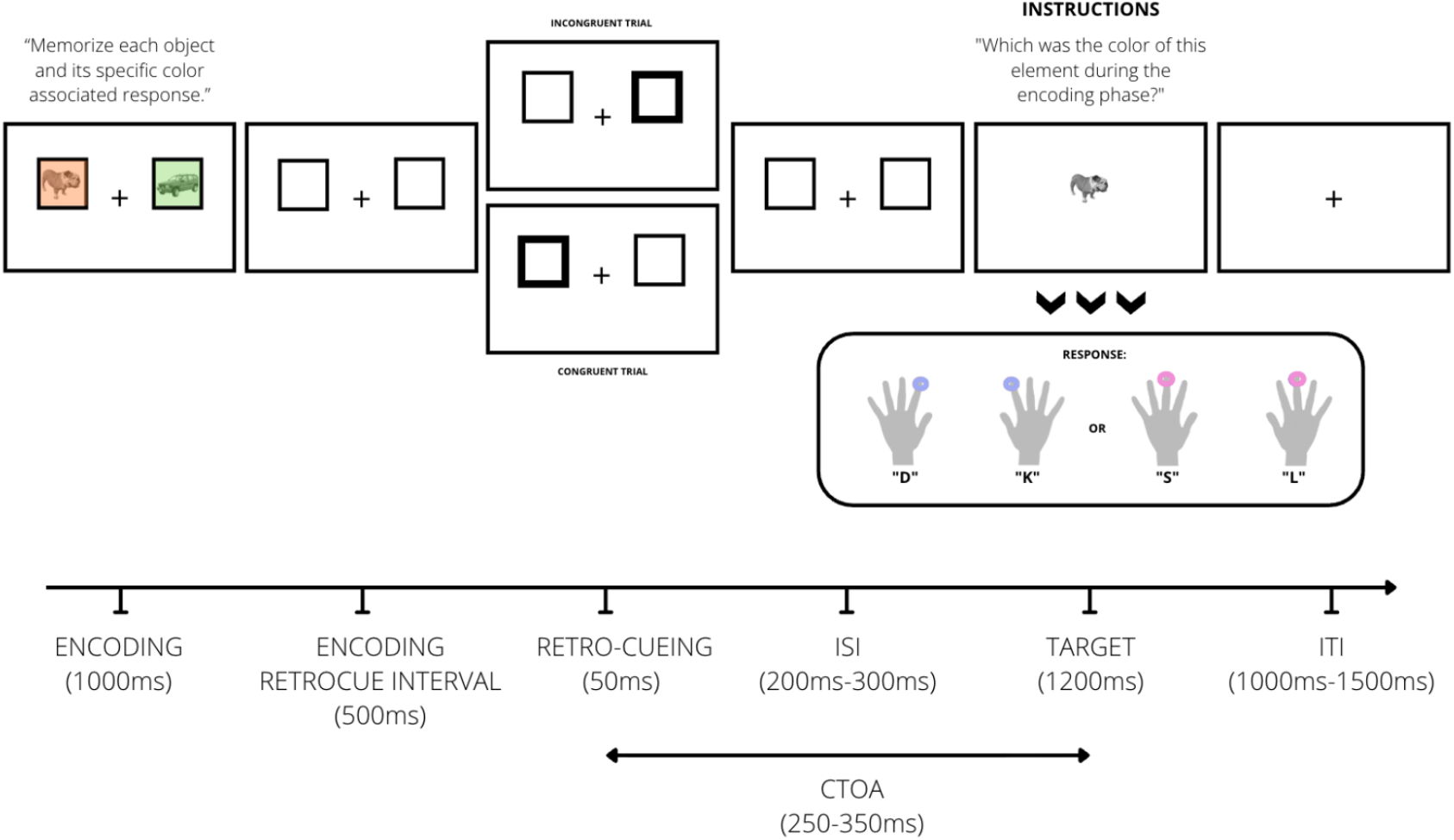
Sequence of events in a given trial. Participants first memorized two distinct stimulus-response (S-R) mappings, each defined by the background color of a placeholder (green or orange). Following the memory encoding phase, an uninformative, non-predictive, exogenous retro-cue flashed in one of the two placeholders, cueing the later probed item or the opposite one (congruent vs. incongruent trials, respectively). This cue was task-irrelevant and instructed to be neglected. Finally, the target stimulus appeared in the center of the screen (rendered in black and white), and participants were required to recall and report the color that was originally associated with the target item through the initially encoded S-R mapping. In a given trial, one of the images was always ‘animate’ while the other was ‘inanimate’. Stimulus animacy was never task-relevant. The retro-cue equally often matched the location of the previous animate or the inanimate image. Note that there was also a neutral block at the end of the experiment in which the cue selected both placeholders, which is not depicted in this figure (see Footnote number 1). *Note*. ISI: inter-stimulus interval. ITI: inter-trial interval. CTOA: cue-target onset asynchrony.

Each participant completed two runs of 299 trials (280 regular, 19 catch), where the retro-cue appeared on a single placeholder (congruent/incongruent trials), and a third run blocked at the end of the experiment of 207 trials (201 regular, 6 catch) in which both placeholders were retro-cued simultaneously (neutral trials^1^), totaling 804 trials. Breaks were allowed midway through each run and between runs. Before the main task, participants completed a practice block of 18 trials (16 regular, 2 catch) without retro-cues. The practice block was repeated until participants achieved at least 85% accuracy. Practice stimuli were not reused in the main task. The full experimental session lasted approximately 170 minutes, including EEG setup and practice phase.

#### Experimental Design

The experiment employed a within-participants design with one factor, cueing, which had two levels: Congruent (the target was the item associated with the previously cued location) and incongruent (the target was the item associated with the opposite location). In addition to standard behavioral measures (accuracy and RTs), we also analyzed participants’ performance using parameters derived from the Hierarchical Drift Diffusion Model (HDDM; Wiecki et al., 2013). EEG data were recorded concurrently during task performance to examine neural correlates of exogenous attentional selection of WM contents.

#### EEG data acquisition and preprocessing

EEG signals were recorded using a 63-channel BrainVision actiCHamp system with active electrodes placed according to the international 10–20 system. Signals were sampled at 500 Hz (sampling interval = 0.002 s) and digitized with a resolution of 0.049 µV. Cz served as the online reference electrode and Fpz as the ground. Impedances were kept below 5 kΩ whenever possible. No online filters were applied during acquisition (DC to 140 Hz; notch filter off). Data were acquired using BrainVision Recorder (v. 1.25.0201).

Offline preprocessing and analyses were conducted in MATLAB using a combination of the FieldTrip toolbox (Oostenveld et al., 2011) and custom scripts developed in the Proactive Brain Lab. The continuous EEG data were segmented into epochs ranging from –200 to +1500 ms relative to retro-cue onset for cue-locked analyses, and from –200 to +2000 ms relative to memory test onset for probe-locked analyses.

EEG data were downsampled from 500 Hz to 250 Hz to reduce computational load. Artifact correction was performed using independent component analysis (ICA), implemented via the runica method in FieldTrip. Components associated with eye blinks and horizontal eye movements were identified and removed based on visual inspection.

Additionally, noisy channels were interpolated manually for specific participants using the average of neighboring electrodes. Trials with excessive variance were identified and rejected using the ft_rejectvisual function with the ‘summary’ method, applied across different frequency bands (4–7 Hz and 8–30 Hz) to aid visual inspection. Artifact rejection was performed blind to experimental condition. On average, the number of remaining trials was 770 [287 congruent, 285 incongruent, and 198 neutral] for cue-locked analyses and 727 trials [267 congruent, 268 incongruent, and 192 neutral] for probe-locked analyses. After excluding neutral trials, 572 trials were considered for the cue-locked analyses and 535 for the probe-locked analyses.

### Statistical analyses

#### Generalized Linear Mixed Models (GLMMs)

Behavioral data were analyzed using two Generalized Linear Mixed Models (GLMMs; see Lo & Andrews, 2015), one for RTs and another for accuracy. In both models, cueing (congruent, incongruent) was included as a fixed effect. To determine the optimal random-effects structure, we compared all possible combinations of random intercepts and slopes for cueing across participants and trials. For each selected model, a Type III Wald chi-square test was conducted. Prior to this, data were filtered to exclude catch trials and for the GLMM on RTs only trials with correct responses were considered. All statistical analyses were performed using RStudio (v2022.02.3) and JASP (v0.14.0.0).

#### Hierarchical Drift Diffusion Model (HDDM)

To further investigate the cognitive mechanisms underlying the cueing effect, we applied HDDM (Wiecki et al., 2013) using Python 3.6 in Spyder. The HDDM framework models two-alternative forced-choice decisions as a process of evidence accumulation toward one of two decision boundaries (Ratcliff & Rouder, 1998), allowing for the estimation of latent parameters that reflect distinct cognitive processes. The model estimates four key parameters: Drift rate (v; speed and quality of evidence accumulation), non-decision time (t^0^; time consumed by processes unrelated to decision-making, e.g., perceptual encoding, motor execution), decision threshold (a; amount of evidence required to make a decision) and starting point (z; initial bias toward one of the decision boundaries).

The hierarchical structure of HDDM allows for the estimation of group-level parameters that inform individual-level estimates, improving stability even with relatively few trials per participant (Lerche et al., 2017). Based on prior work using the same task (Fuentes-Guerra et al., 2025a), trials with RTs below 200 ms were excluded. Only regular (non-catch) trials were included in the analysis. To account for potential outliers, we used the p_outlier parameter in HDDM to specify a mixture model assuming 5% of trials originated from a uniform distribution. Model estimation was performed using Markov Chain Monte Carlo (MCMC) sampling with 10,000 samples per chain (four chains). The first 1,000 samples were discarded as burn-in, and every second sample was removed to reduce autocorrelation. Convergence was assessed using the R-hat statistic, with all values below 1.05, indicating satisfactory convergence (Makowski et al., 2019).

We compared three models differing in whether cueing modulated both v and t^0^, only v, or only t^0^. The differences in Deviance Information Criterion (DIC) between models exceeded the conventional threshold of 5 (Cain & Zhang, 2019). Therefore, we selected the second model as the most parsimonious model for interpretation (see Table 1) since it was closest to zero. This model replicated previous findings (Fuentes-Guerra et al., 2025a), showing that cueing primarily affected the drift rate, suggesting that attentional cueing modulates the quality of evidence entering the decision process.

**Table 1.** Model fitting between the three proposed drift diffusion models.

| Model # | Cueing | DIC |
| --- | --- | --- |
| 1 | $v, t_0$ | -4078.29 |
| 2 | $v$ | -3806.83 |
| 3 | $t_0$ | -4079.76 |

*Note.* $v$ : drift rate; $t_0$ : non-decision time; $a$ : decision threshold; DIC: Deviance Information Criterion

#### EEG Analyses

#### Multivariate Pattern Analyses (MVPA)

To assess whether, at cue presentation, the EEG signal over posterior electrodes contained content-specific information about the memorized item, we implemented a time-resolved multivariate decoding approach using the Mahalanobis continuous distance (Mahalanobis, 1936; De Maesschalck et al., 2000). Analyses focused on posterior channels of interest (Pz, P3, P5, P7, P4, P6, P8, Oz, O1, O2, POz, PO3, PO4, PO7, PO8; See Fig. 3A), based on previous literature in visuospatial attention (Wang, et al., 2023; Liu & van Ede, 2025) and sensory “pinging” (Wolff, et al., 2015, 2017). Trials were categorized based on animacy (animate vs. inanimate) and cue position (left vs. right). To ensure balanced conditions, we randomly sampled an equal number of trials per condition (cued animate left/right and cued inanimate left/right). Note that “cued” refers to the item selected by the retro-cue at the time of its presentation, regardless of its eventual congruency, which cannot be determined until the probe appears.

For decoding, we used a leave-one-trial-out cross-validation procedure: each trial was iteratively treated as a test trial, and all remaining trials served as the training set. For each time point, we computed the similarity between the test trial and the average pattern of matching trials (same class) and non-matching trials (different class). The Mahalanobis continuous distance, a (dis)similarity measure, was computed as the difference between the distance to the non-matching class and the distance to the matching class, such that positive values indicate greater similarity to the matching class (Wolff et al., 2015, 2017). This procedure was repeated for all trials and time points, resulting in a time-resolved decoding accuracy measure. The main goal of this analysis was to test whether, for cued items, the animacy of the previously presented item at that location could be decoded from the EEG signal. This would serve as evidence for retro-cue driven reactivation of the spatially matching WM content.

Additionally, to fully identify the spatial origin of neural patterns that support the discrimination between cued animate and inanimate objects held in WM, we implemented a searchlight decoding approach (as in e.g., van Ede et al, 2019), including the whole electrode set. For each electrode, a local cluster was defined comprising the electrode and its spatial neighbors, based on the EasyCap layout. EEG data from each trial were extracted for the selected cluster. Then, the same decoding analysis as above was iteratively run, but for each electrode and its adjacent neighbors, to yield a topography.

#### Event Related Potentials (ERPs)

To examine neural responses at probe presentation, Event Related Potentials (ERPs) were computed over the same posterior electrodes used for the MVPA analysis for both congruent and incongruent trials (See Fig. 4A). Data were epoched from −200 to 1000 ms relative to probe onset, baseline-corrected using the −250 to 0 ms interval, and low-pass filtered at 30 Hz. Trials with artifacts were excluded. Scalp topographies were generated using the FieldTrip function ft_topoplotER.

#### Cluster-Based Permutation Tests on EEG data

To statistically assess the effects of exogenous cueing on the representational and temporal dynamics of the EEG signal, we employed a cluster-based permutation approach implemented via FieldTrip’s ft_timelockstatistics. This non-parametric method is well-suited for evaluating spatiotemporal consistency in EEG data and controls the multiple comparisons problem by generating a permutation distribution of the largest cluster statistic under the null hypothesis. In our implementation, clusters were formed by grouping temporally adjacent data points with the same sign that exceeded a predefined significance threshold in a mass-univariate two-sided dependent-samples t-test (α = 0.05). Cluster size was defined as the sum of all t-values within each cluster. We used 10,000 permutations to construct the null distribution of the maximum cluster statistic. The function assumes a within-participant design (dependent samples) and uses FieldTrip’s Monte Carlo method for cluster correction when statMethod = ‘montecarlo’.

This procedure was applied to two main analyses: (1) to assess the strength of the decoding signal distinguishing cued animate from cued inanimate items at the time of cue presentation, and (2) to evaluate the significance of the congruency effect in the ERPs following probe onset.

## Results

In the following, we first confirm that exogenous retro-cues facilitate performance to spatially matching vs. mismatching WM contents, before turning to our main EEG analyses targeting retro-cue driven reactivation of WM content and neural processing of the WM probe.

### Retrospective selection of spatially matching memories affects performance

To examine the effect of cueing (difference between congruent and incongruent WM probes; see Fig. 2) on RTs, we compared five GLMMs with varying random-effects structures. The model with the lowest Akaike Information Criterion (AIC = 162,799.36) and Bayesian Information Criterion (BIC = 162,866.42) was selected for further analysis. This model included cueing as a fixed effect and allowed both the baseline RTs (intercepts) and the effect of cueing (slopes) to vary across participants and trials. A Type III Wald chi-square test revealed a significant main effect of cueing on RTs, χ^2^(1, N = 25) = 31.21, p <.001, indicating that participants responded significantly faster on congruent trials (M = 678 ms, SD = 99 ms) compared to incongruent trials (M = 715 ms, SD = 98 ms) (See Fig. 2A, left upper panel). The average cueing effect (congruent – incongruent trials) was −37 ms (SE = 4.28 ms) (see Fig. 2B, right upper panel).

**Fig. 2.**
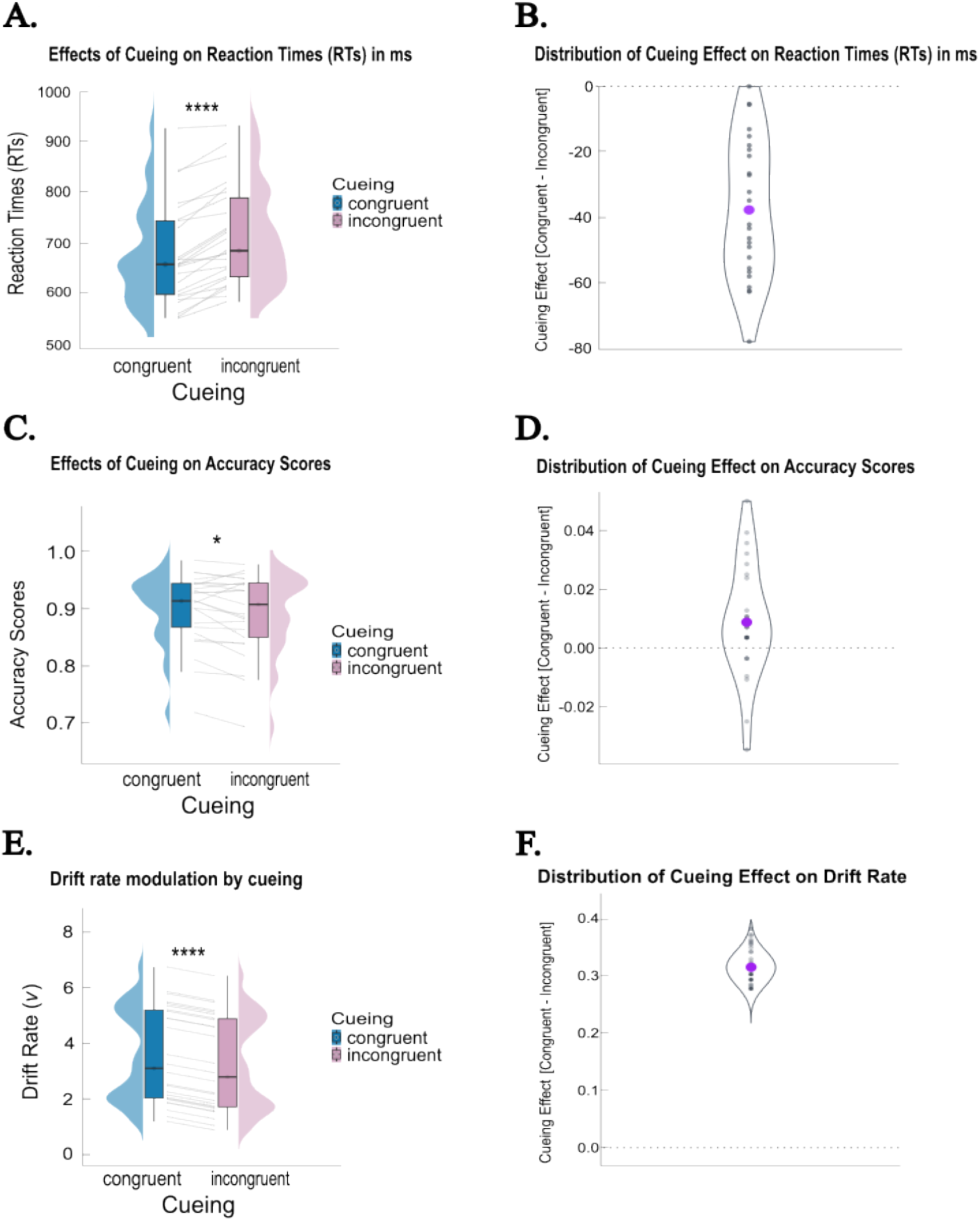
Behavioral results. **A**. Effects of cueing on Reaction Times (RTs). *Note*. The maximum and minimum Reaction Times’ (RTs) values are represented in the whiskers of the box plots. The Interquartile range (IQR) is displayed in the boxes by portraying the lower quartile, median and upper quartile. The half-violin plots represent the distribution of RTs across conditions. The grey lines represent individual participants. The asterisks represent statistical significance. **B**. Cue-congruency effects on RTs quantified as the differences in RT following congruent minus incongruent WM probes. *Note*. The larger purple dot represents the average cueing effect. **C**. Effects of cueing on Accuracy Scores. **D**. Cue-congruency effects on accuracy scores. **E**. Drift rate modulation by cueing. **F**. Cue-congruency effects on drift-rate quantified as the difference in drift rate (v) between congruent and incongruent WM probes. *Note*. The violin plot represents the posterior distribution of the group-level effect estimated via the HDDM, and the dots represent samples from this distribution. The larger purple dot represents the mean of the posterior.

**Fig. 3.**
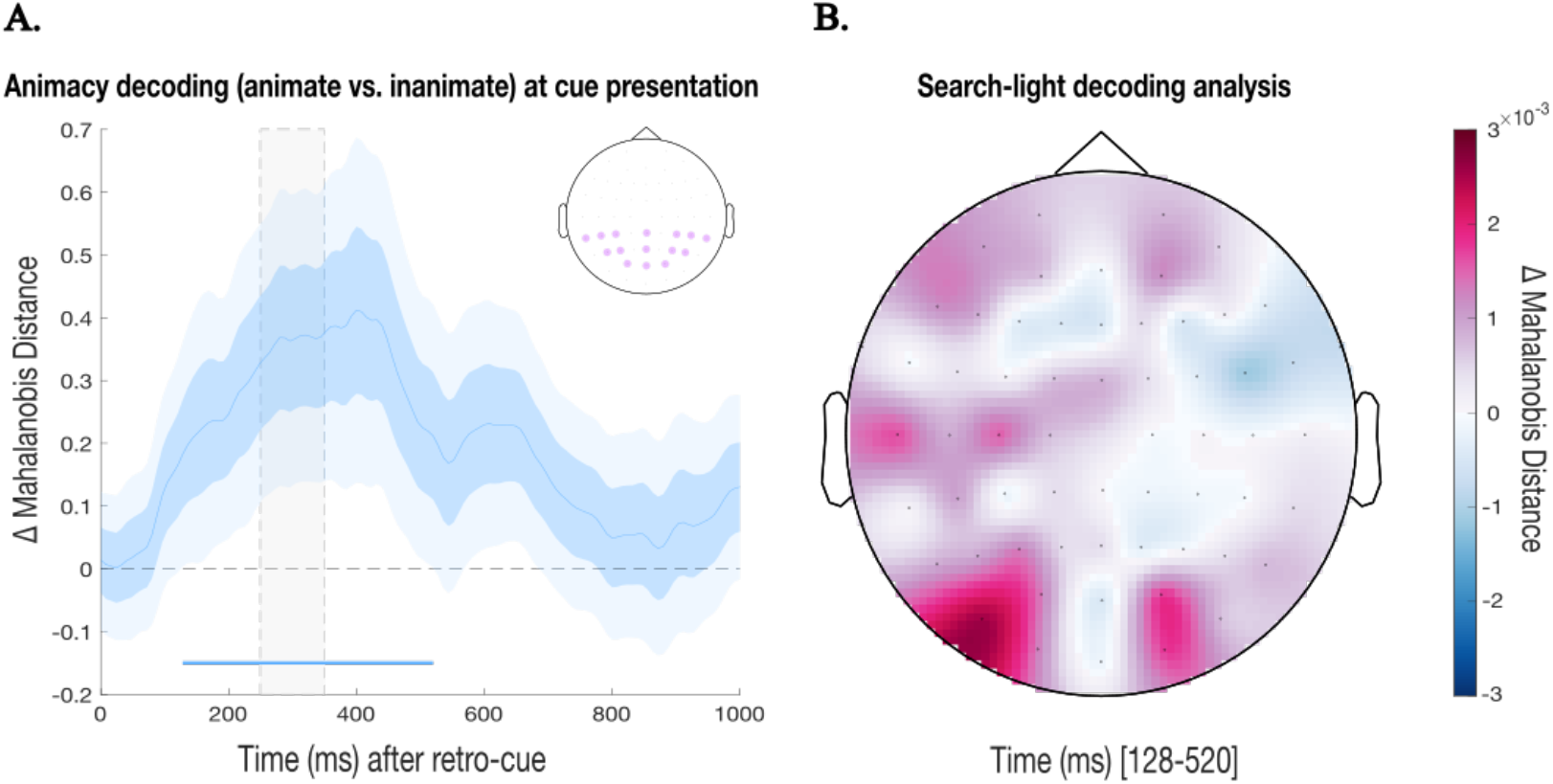
EEG decoding results. **A**. Time course of Mahalanobis animacy decoding (cued animate vs. cued inanimate) over posterior electrodes following cue presentation (t=0). Decoding was performed using a leave-one-out approach and quantified as the differences in Mahalanobis distance between a left-out test train and the average of all other trials (training trials) with matching vs. mismatching animacy labels (mismatching minus matching, such that a small distance to matching trials yields positive decoding). *Note*. The vertical grey, dashed, shaded region represents de jittered interval when the probe appears (t=250-350ms). Darker blue shaded regions represent standard errors (SE) and lighter blue shaded regions represent the 95% confidence intervals (CI). The horizontal blue bar (near the x-axis) indicates the time window of the significant positive cluster (128–520 ms, p <.05) identified by a cluster-based permutation test. The embedded topography on the upper right corner shows the selected electrodes for the MVPA analyses, highlighted in shaded purple. **B**. Topographic distribution of Mahalanobis continuous distance for animacy decoding during the significant cluster (128–504 ms after cue onset).

**Fig. 4.**
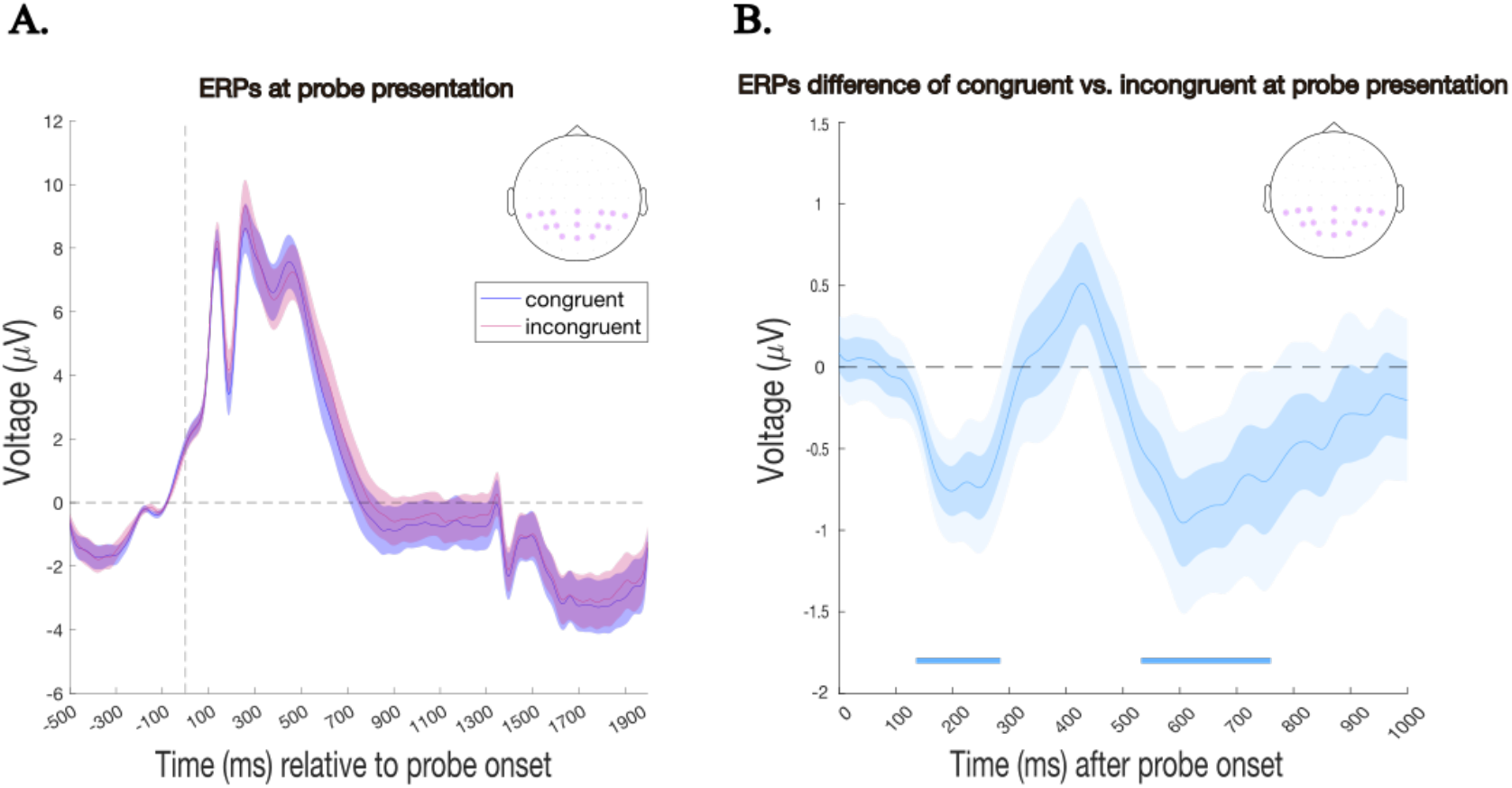
**A**. Event-related potentials (ERPs) recorded over posterior electrodes following probe onset, separately for congruent (blue) and incongruent (red) conditions. *Note*. The vertical dashed line represents probe onset (t=0). Darker shaded regions represent standard errors (SE). **B**. Difference wave (congruent minus incongruent) for ERPs over posterior electrodes following probe onset (t=0). The embedded topography on the upper right corner shows the selected electrodes for the ERP analyses, highlighted in shaded purple. *Note*. Horizontal blue bars mark time windows with significant differences (cluster-based permutation test, p <.05). Two clusters emerged: an early effect (136–284 ms) and a later effect (536–756 ms) both reflecting relatively greater negativity (smaller positivity) for congruent probes compared to incongruent probes. Darker shaded regions represent standard errors (SE) and lighter shaded regions represent 95% confidence intervals (CI).

Regarding accuracy, the selected model with the lowest AIC (8965.16) and BIC (8995.41) included cueing as a fixed effect and random intercepts for both participants and trials to account for variability in baseline accuracy. A Type III Wald chi-square test provided a significant main effect of cueing on accuracy scores, χ^2^(1, N = 25) = 219.66, p <.001, indicating that participants responded more accurately on congruent trials (M =.90, SD =.06) compared to incongruent trials (M =.89, SD =.07) (See Fig. 2C, left middle panel). The average cueing effect (congruent – incongruent trials) was.008 (SE = 0.004) (see Fig. 2D, right middle panel).

Last, using HDDM, RTs and accuracy were jointly modeled. After evaluating model fit, the best-fitting model was the one in which cueing only modulated drift rate (DIC = −3806.83). We found a main effect of cueing (b = −0.315, 95% HDI = [−0.369, −0.264], pd = 100%), replicating our previous findings (Fuentes-Guerra et al., 2025a), and suggesting that evidence accumulation was faster in congruent compared to incongruent trials (Fig. 2E). The average cueing effect (congruent – incongruent trials) was 0.313 (Posterior SD = 0.028) (see Fig. 2F).

### Exogenous retro-cues selectively revive neural traces of spatially matching WM content

We now turn to our main analyses where we performed MVPA on posterior EEG electrodes to assess whether, following retro-cue presentation, content-specific information about the memorized item emerged in the EEG signal before probe onset. To this end, we decoded the ‘animacy’ of the memorized object on the side of the spatial retro-cue. This showed that the animacy of the cued item could be significantly decoded above-chance after cue presentation (see Fig. 3A). That is, an exogenous non-predictive and task-irrelevant spatial retro-cue selectively and automatically revived the category (animacy) of spatially matching WM content. Note how this was the case even though the cue itself was purely spatial with no animacy information whatsoever. The cluster-based permutation test revealed a significant positive cluster from 128 to 520 ms after cue onset, indicating above-chance animacy decoding over posterior electrodes following cue presentation. This shows reliable differentiation between cued animate and inanimate stimuli during this interval in response to the spatial retro-cue, when no items were presented on the screen apart from the purely spatial cue, and persisted through the onset of the probe at 250–350 ms. The searchlight decoding analysis corroborated a predominantly posterior contribution to this decoding (see Fig. 3B), consistent with the reactivation of category-specific animacy information in a “visual” format.

### Congruency effects emerge at early stages following WM probe presentation

We finally investigated neural responses to the WM probes at the end of the WM delay. Following uninformative cues, target probes could be congruent if they matched the previously selected item or incongruent if the probe tested the other WM item. If cues affect WM content already at the cue stage (as implied by the previous analysis), then we might also expect differential processes at the later probe (test) stage when probing the same/cued (congruent) vs the other/uncued (incongruent) memory item. Thus, to examine whether probe congruency influenced neural responses, we analyzed ERPs over posterior electrodes following probe onset. Figure 4A shows the grand-average ERPs for congruent and incongruent probes across time. Both conditions exhibited a pronounced positive deflection shortly after probe onset, followed by a gradual return toward baseline. However, visual inspection hinted at a more negative-going waveform for congruent relative to incongruent probes during early post-probe intervals. To quantify these differences, we computed the ERP difference wave (congruent minus incongruent) and assessed statistical significance across time (See Fig. 4B). Two significant clusters emerged: an early cluster between 136–284 ms and a later cluster between 536–756 ms post-probe. Both clusters indicate relatively greater negativity (smaller positivity) for congruent probes compared to incongruent probes.

Consistent with the previous decoding findings, these complementary probe-locked ERP findings show that the preceding exogenous retro-cue affects probe processing early on, consistent with exogenously cued (congruent) vs. uncued (incongruent) items being in a distinct state when the probe comes in. At the same time, congruency between the probed memory content and previously cued content additionally appears to affect a subsequent later second stage, possibly associated with decision-making and response execution (approximately around the average RTs).

## Discussion

This study investigated whether and how exogenous, non-predictive and task-irrelevant retro-cues automatically reactivate visual WM contents. To this end, EEG signals were recorded to perform multivariate decoding and ERP analyses, aimed at evaluating the processes occurring at cue presentation and at probe, respectively. Three main findings emerge from our data, which we discuss in turn.

First, replicating previous behavioral work (Cipriani et al., 2024; Ding et al., 2024; Fuentes-Guerra et al., 2025a, Fuentes-Guerra et al., 2025b; Han et al., 2023; van Ede et al., 2020), we show that non-predictive exogenous retro-cues can drive attention to complex WM contents. This effect is evident in both RTs and accuracy, with faster and more accurate responses in congruent trials compared to incongruent trials, mimicking the pattern of external exogenous attentional selection (Posner, 1980; Chica et al., 2013). Drift diffusion modelling further indicates that this retro-cueing effect on performance is primarily driven by faster evidence accumulation in congruent compared to incongruent trials, as only drift rate was modulated by exogenous retro-cues. This pattern is broadly consistent with ideas proposed by the matched filter hypothesis (Hayden & Gallant, 2013; Myers et al., 2015; Muhle-Karbe et al., 2021) and the protection-from-interference hypothesis (Makovski et al., 2008), although our study was not designed to adjudicate between these accounts. Both frameworks offer plausible interpretations: the former emphasizes improved efficiency in extracting relevant information, while the latter highlights shielding the cued item from interference (for a review, see Fuentes-Guerra et al., 2025c). In sum, behavioral results provide evidence that exogenous retrospective selection of spatially matching memories improves performance via enhanced evidence accumulation, which seems to protect the representations and/or enhance how efficiently relevant information is extracted to come up with a decision.

Second, our study uniquely demonstrates that exogenous retro-cueing can selectively revive neural traces of matching WM content at cue presentation. This extends previous evidence from global sensory “pinging” approaches used to probe activity-silent WM traces (Barbosa et al., 2021; Karabay et al., 2025; Wolff et al., 2015, 2017; Yang et al., 2025; Zhang & Luo, 2023). Earlier studies showed that memory-item-specific information could be decoded from the impulse response even in the absence of sustained delay activity or attention. However, this reactivation was not selective: the impulse response reflected both attended and unattended items in WM. This likely stems from their perturbation method, which used a high-contrast, task-irrelevant visual input composed of three circles simultaneously occupying the positions of the two memorized gratings and the fixation point, thereby producing a global rather than item-specific reactivation. Within this global perturbation approach, Yang and collaborators (2025) highlighted the role of space matching in neural reactivation by presenting either three vertical circles (spatially non-matching) or three horizontal circles (spatially-matching) as “pings”. They found that spatial-nonmatching pings transiently reactivated activity-silent WM without altering original WM representations or recall performance, whereas spatial-matching pings produced more durable reactivation and reorganized WM information by reducing representations’ dynamics. Notably, only the reactivation strength of spatial-matching pings correlated with recall. Building on this idea, our experiment went one step further: the retro-cue always selected one of the two items, allowing discrimination of the cued item’s animacy (animate vs. inanimate), thus providing direct evidence for content-specific reactivation of the spatially matching item. Note how the cue itself never contained any information regarding animacy, such that the cue-induced animacy decoding must reflected activation of the spatially matching visual object held in WM.

This finding parallels established frameworks from long-term memory and sleep research on targeted memory reactivation (TMR), in which external cues are used to selectively trigger the reactivation of latent memory traces (Rasch & Born, 2007; Lewis & Bendor, 2019; Herszage & Censor, 2017). In this approach, sensory cues such as odors (Rasch et al., 2007) or sounds (Rudoy et al., 2009) are paired with information during wakefulness and later re-presented during subsequent sleep (Lewis & Bendor, 2019). This procedure enhances the reactivation of cued memories, and such enhancement predicts later memory performance (for a review, see Cellini & Cappuzo, 2018). The replay of information appears to rely on cortical inputs to the hippocampus (Buzsaki, 1989; Ji & Wilson, 2007; Issa & Wang, 2008). We now show this in the context of WM, where such a phenomenon has remained surprisingly underexplored.

We also note how our task involved judging colour, not animacy. Yet, our data show how animacy was automatically decoded at the time of retro-cue presentation. This suggests that exogenous retro-cues not only select the task-relevant features of the memory object itself but also the entire “event” of which it is a part. This supports the predictions posed by the Theory of Event Coding (TEC; Hommel, 2019), which have been reassessed (Janczyk et la., 2022), and built upon in the Binding and Retrieval in Action Control (BRAC) Framework (Frings et al., 2020). According to BRAC, stimulus features, responses, environmental context, and their subsequent effects are integrated into a “mental representation” that includes all the elements related to a specific event (Frings et al., 2020). Such mental representation is referred to as an event file, as coined by Hommel, Müsseler, Aschersleben, and Prinz (2001); Hommel (2019). Importantly, it is assumed that the latter repetition of any of these elements triggers the retrieval of the previous event-file as a whole, comprising codes of the same features, which can impact current performance (Frings et al., 2020). In this case, a spatial retro-cue reactivated the event previously contained in that specific location, as evidence from a reactivation of the event’s animacy, even though animacy was technically irrelevant to the memory task.

Third, during cue presentation, only one of the two items was selected, yet this selection was not predictive of which item would appear as the probe at the end of the WM delay. Consequently, trial type (congruent or incongruent) was determined solely by the identity of the final probe. A trial was classified as congruent if the probe matched the previously selected item, or incongruent if it was the unselected item. Thus, the congruency manipulation was only revealed at the end of the trial upon probe presentation. If cues bring matching memory contents into a different ‘state’, we may nevertheless expect a difference in processing of the probe when it probes the previously cued (congruent) vs the other (incongruent) memory item. To assess this, ERP analyses time-locked to the probe were performed. These analyses revealed that congruency effects emerged at early sensory/visual stages following probe presentation (Herrmann, & Knight, 2001). Specifically, already from approximately 136 to 284 milliseconds after probe onset, a relatively greater negative (less positive) deflection was observed for congruent trials compared to incongruent ones. This pattern may be interpreted as a significant modulation in early components such as P1 and N1, which are well-established early markers of enhanced sensory processing. These rapid effects suggest that exogenous attention during WM may boost perceptual encoding of congruent probes prior to higher-level decision processes (Hillyard & Mangun, 1987; Luck, 1995; Vogel & Luck, 2000). In addition, we also observed a second, later effect during probe processing. This late modulation likely signals additional processes, such as later decision-making and response preparation or execution, which is consistent with the RT differences we recorded between the congruent and incongruent trials.

In the present study, after examining category-based representational decoding patterns following retro-cue presentation, our findings demonstrate that uninformative, non-predictive exogenous spatial retro-cues presented during the WM delay can act as selective “pings”, reactivating spatially matching WM content. Considering the TEC and BRAC frameworks (Hommel, 2019; Frings et al., 2020), future work could test whether any feature can act as a selective “ping” (e.g., a color-based retro-cue, cf. van Ede et al. 2020), or whether this process of full-event reactivation is idiosyncratic to spatial exogenous selection, which is characterized by its automaticity (Chica et al., 2013). Moreover, future work should also assess whether such automatic exogenous selection produces longer-lasting effects, affecting also long-term memory representations. Confirmation of such findings would have critical implications for how the external environment can involuntarily penetrate our internal cognitive workspace and influence subsequent actions. Finally, it would be important to dissect the neural circuitry responsible for the here reported effects, such as the potential contribution of dorsal and/or ventral attention networks (Chica et al., 2013; Corbetta et al., 2008) that have been implicated in exogenous attention and WM (Yoon et al., 2008; Zhou et al., 2022), the oculomotor system that has been implicated in spatial attention shifts in perception and WM (Awh et al., 2006; Awh & Jonides, 2001; de Vries & van Ede, 2024; Rolfs, 2009; Liu et al., 2024), and/or the type of hippocampal-cortical loops that have been implicated in targeted memory reactivation in the context of sleep (Buzsaki, 1989; Ji & Wilson, 2007; Issa & Wang, 2008; Rothschild, 2019).

## Conflict of interest statement

“The authors declare no competing financial interests.”

## Acknowledgments

This work was supported by the Spanish Ministry of Science and Innovation and the State Research Agency [research Project PID2020-116342GA-I00 to EMA and CGG, PID2023-149428NB-I00 to CGG, and PID2024-157672NB-I00 to EMA, funded MICIU/AEI/10.13039/501100011033/FEDER, EU]. CGG was also supported by Grant Ramón y Cajal RYC2021-033536-I funded by MCIN/AEI/10.13039/501100011033 and by the European Union Next Generation EU/PRTR. FvE was supported by an ERC Starting Grant from the European Research Council (MEMTICIPATION, 850636) and an NWO Vidi Grant by the Dutch Research Council (grant number 14721).

AFG was supported by Grant PRE2021-100351, funded by MCIN/AEI/10.13039/501100011033. Additionally, this publication was funded by the European Social Fund Plus ESF+, CEX2023-001312-M by MCIN/AEI/10.13039/501100011033 and by the UCE-PP2023-11 by the University of Granada as an Excellence Unit Program. This work is part of the doctoral thesis of AFG, under the supervision of EMA and CGG.

## CRediT authorship contribution statement

**Á. Fuentes-Guerra**: Conceptualization, Methodology, Software, Validation, Formal analysis, Investigation, Data Curation, Writing – original draft, Writing – review & editing, Visualization, Project administration.

**E. Martín-Arévalo:** Writing – review & editing, Validation, Supervision, Resources, Methodology, Funding acquisition, Conceptualization.

**F. van Ede:** Writing – review & editing, Validation, Supervision, Software, Formal analysis, Resources, Conceptualization.

**C. González-García:** Writing – review & editing, Validation, Supervision, Software, Resources, Methodology, Funding acquisition, Conceptualization.

## Declaration of generative AI and AI-assisted technologies in the writing process

During the preparation of this work the author(s) used ChatGPT-4 (GPT-4-turbo), accessed via Microsoft Copilot, to assist with English language correction. The author(s) reviewed and edited the content as necessary and take full responsibility for the final version of the manuscript.

## Footnotes

1 Neutral trials in exogenous attention paradigms are hard to interpret. They should provide a baseline, but often responses are unexpectedly faster or slower than cued and uncued trials. This may reflect alerting effects, visual properties, trial frequency, or inconsistent strategies, such as participants might diffuse attention or adopt idiosyncratic strategies. Because of this, neutral trials can be problematic, and their interpretation requires caution (Chica et al., 2014). Following this reasoning, neutral trials were presented at the end of the experiment to minimize interference with exogenous cueing effects, in case they would be of interest for future re-analyses of the dataset, while not being central for the current study. However, we note that this design choice introduces certain limitations as well: behavioral performance may improve due to learning effects, and EEG signals tend to drift and become noisier over time. Therefore, neutral trials were not included in the main analyses here but are available in the online repository (https://openneuro.org/datasets/ds007180; https://osf.io/e3gxj/overview?view_only=ad6c91039e5f40bc9bebd8d998a27812).

## Notes

https://openneuro.org/datasets/ds007180

### Competing Interest Statement

The authors have declared no competing interest.

